# Leveraging omic features with F3UTER enables identification of unannotated 3’UTRs for synaptic genes

**DOI:** 10.1101/2021.03.08.434412

**Authors:** Siddharth Sethi, David Zhang, Sebastian Guelfi, Zhongbo Chen, Sonia Garcia-Ruiz, Mina Ryten, Harpreet Saini, Juan A. Botia

## Abstract

There is growing evidence for the importance of 3’ untranslated region (3’UTR) dependent regulatory processes. However, our current human 3’UTR catalogue is incomplete. Here, we developed a machine learning-based framework, leveraging both genomic and tissue-specific transcriptomic features to predict previously unannotated 3’UTRs. We identify unannotated 3’UTRs associated with 1,513 genes across 39 human tissues, with the greatest abundance found in brain. These unannotated 3’UTRs were significantly enriched for RNA binding protein (RBP) motifs and exhibited high human lineage-specificity. We found that brain-specific unannotated 3’UTRs were enriched for the binding motifs of important neuronal RBPs such as *TARDBP* and *RBFOX1*, and their associated genes were involved in synaptic function and brain-related disorders. Our data is shared through an online resource F3UTER (https://astx.shinyapps.io/F3UTER/). Overall, our data improves 3’UTR annotation and provides novel insights into the mRNA-RBP interactome in the human brain, with implications for our understanding of neurological and neurodevelopmental diseases.

## Introduction

The 3’UTRs of protein-coding messenger RNAs (mRNAs) play a crucial role in regulating gene expression at the post-transcriptional level. They do so by providing binding sites for *trans* factors such as RBPs and microRNAs, which affect mRNA fate by modulating subcellular localisation, stability and translation [1, 2]. There is evidence to suggest that these RNA-based regulatory processes may be particularly important in large, polarised cells such as neurons. Recent studies have shown that transcripts which are highly expressed in neurons have both significantly longer 3’UTRs and higher 3’UTR diversity [3, 4]. Furthermore, it has been shown that thousands of mRNA transcripts localise within subcellular compartments of neurons and undergo regulated local translation, allowing neurons to rapidly react to local extracellular stimuli [4–7]. Thus, there has been growing interest in the impact of 3’UTR usage on neuronal function in health and disease.

However, despite on-going efforts to identify and characterise 3’UTRs in the human genome [8–11], there is evidence to suggest that our current catalogue is incomplete [3, 12–14]. Large-scale 3’end RNA-sequencing (RNA-seq) has identified a large number of novel polyadenylation (poly(A)) sites, many of which are located outside of annotated exons [12, 13]. These insights are complemented by an increasing recognition of the functional importance of transcriptional activity outside of known exons, particularly in human brain tissues [15–17]. This raises the possibility of developing new approaches for 3’UTR identification seeded from RNA-seq data analyses, an area that has not been fully explored, in large part due to the limited availability of data and appropriate tools.

In this study, we present a machine learning-based framework, named F3UTER, which leverages both genomic and tissue-specific transcriptomic features. We apply F3UTER to RNA-seq data from Genotype-Tissue Expression Consortium (GTEx) to predict hundreds of unannotated 3’UTRs across a wide range of human tissues, with the highest prevalence discovered in brain. We provide evidence to suggest that these unannotated 3’UTR sequences are functionally significant and have higher human lineage specificity than expected by chance. More specifically, we found brain-specific unannotated 3’UTRs were enriched for genes involved in synaptic function and interact with neuronal RBPs implicated in neurodegenerative and neuropsychiatric disorders. We release our data in an online platform, F3UTER (https://astx.shinyapps.io/F3UTER/), which can be queried to visualise unannotated 3’UTR predictions and the omic features used to predict them.

## Results

### Annotation-independent expression analysis suggests the existence of unannotated 3’UTRs in the human brain

There is growing evidence to suggest that the annotation of the human brain transcriptome is incomplete and disproportionately so when compared to other human tissues [15–17]. We hypothesised that this difference may in part be attributed to an increased number of unannotated 3’UTRs in human brain. To investigate this possibility, we analysed unannotated expressed regions of the genome (termed ERs) as previously reported by Zhang and colleagues [15]. These ERs were identified through annotation-independent expression analysis of RNA-seq data generated by GTEx with ER calling performed separately for 39 human tissues, including 11 non-redundant human brain regions. We focused on the subset of ERs most likely to be 3’UTRs, namely intergenic ERs which lie within 10 kb of a protein-coding gene (**Methods**). We found that these intergenic ERs were significantly higher in number (*p =* 1.66 × 10^−6^, Wilcoxon Rank Sum Test) and total genomic space (*p =* 2.39 × 10^−9^, Wilcoxon Rank Sum Test) in brain compared to non-brain tissues (**Figure 1a**). Furthermore, we discovered that intergenic ERs were significantly more likely to be located at 3’- rather than 5’-ends of their related protein-coding genes (*p =* 2.08 × 10^−14^, Wilcoxon Rank Sum Test) (**Figure 1b**), suggesting that a proportion of ERs detected in human brain could represent unannotated 3’UTRs.

**Figure 1.**
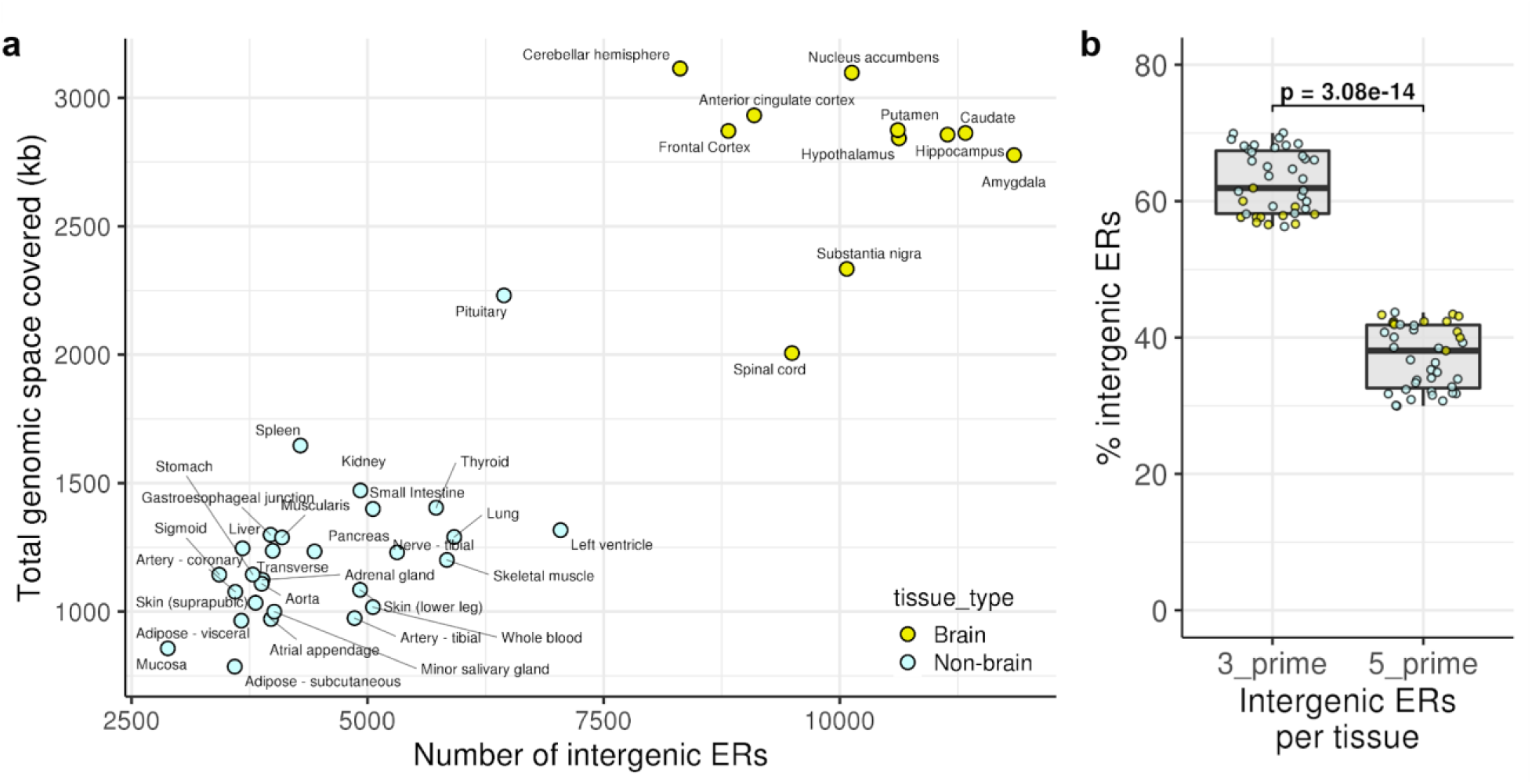
Enrichment of intergenic ERs across 39 GTEx tissues. **(a)** Scatter plot showing the number of intergenic ERs and their total genomic space covered in 39 human tissues. **(b)** Enrichment of intergenic ERs grouped by location with respect to their associated protein-coding gene. Each data point in the box plot represents the proportion of total intergenic ERs in a tissue. p: p-value calculated using Wilcoxon Rank Sum Test.

### Differentiating 3’UTRs from other expressed genomic elements is challenging

Given that existing studies indicate high levels of transcriptional noise and non-coding RNA expression in intergenic regions [18–21], only some intergenic ERs are likely to be generated by unannotated 3’UTRs. This prompted us to develop a method to distinguish 3’UTRs from other transcribed genomic elements (non-3’UTRs) using short-read RNA-seq data. To achieve this aim, we first constructed a training set of known 3’UTRs (positive examples) and non-3’UTRs (negative examples) from Ensembl human genome annotation (v94). We obtained 17,719 3’UTRs and a total of 162,249 non-3’UTRs, consisting of five genomic classes: 21,798 5’UTRs, 130,768 internal coding exons (ICE), 3,718 long non-coding RNAs (lncRNAs), 3,819 non-coding RNAs (ncRNAs) and 2,146 pseudogenes (**Methods**). For each of the positive and negative examples, we constructed a set of 41 informative omic features, which were broadly categorised as either genomic or transcriptomic in nature. Features calculated from genomic data included poly(A) signal (PAS) occurrence, DNA sequence conservation, mono-/di-nucleotide frequency, transposon occurrence and DNA structural properties. Features calculated from transcriptomic data included entropy efficiency of the mapped reads (EE) and percentage difference between the reads mapped at the boundaries (PD) (**Methods**). To gain a better understanding of these features, we performed a univariate analysis to individually inspect the relationship between each feature and the genomic classes in our training dataset (i.e. 3’UTRs and all types of non-3’UTRs). Overall, while the genomic and transcriptomic features used had overlapping distributions amongst some genomic classes, each feature was significantly different when compared across all the genomic classes (*p* < 2.2 × 10^−16^, Kruskal-Wallis Test and proportion Z-Test, **Supplementary Figure S1**). This suggested that the features selected could be used to distinguish 3’UTRs from other genomic elements.

To further investigate this for all 41 features across all six genomic classes, we applied a uniform manifold approximation and projection (UMAP) [22] for dimensionality reduction into a 2D projection space. We found that while most 3’UTRs clustered separately from other classes within that space, some of them highly overlapped with other genomic classes such, as lncRNAs, ICEs and 5’UTRs (**Figure 2a, Supplementary Figure S2**). These findings suggested that many unannotated 3’UTRs would be difficult to identify, and thus, may require an advanced classification approach based on machine learning to accurately distinguish them from other genomic elements.

**Figure 2.**
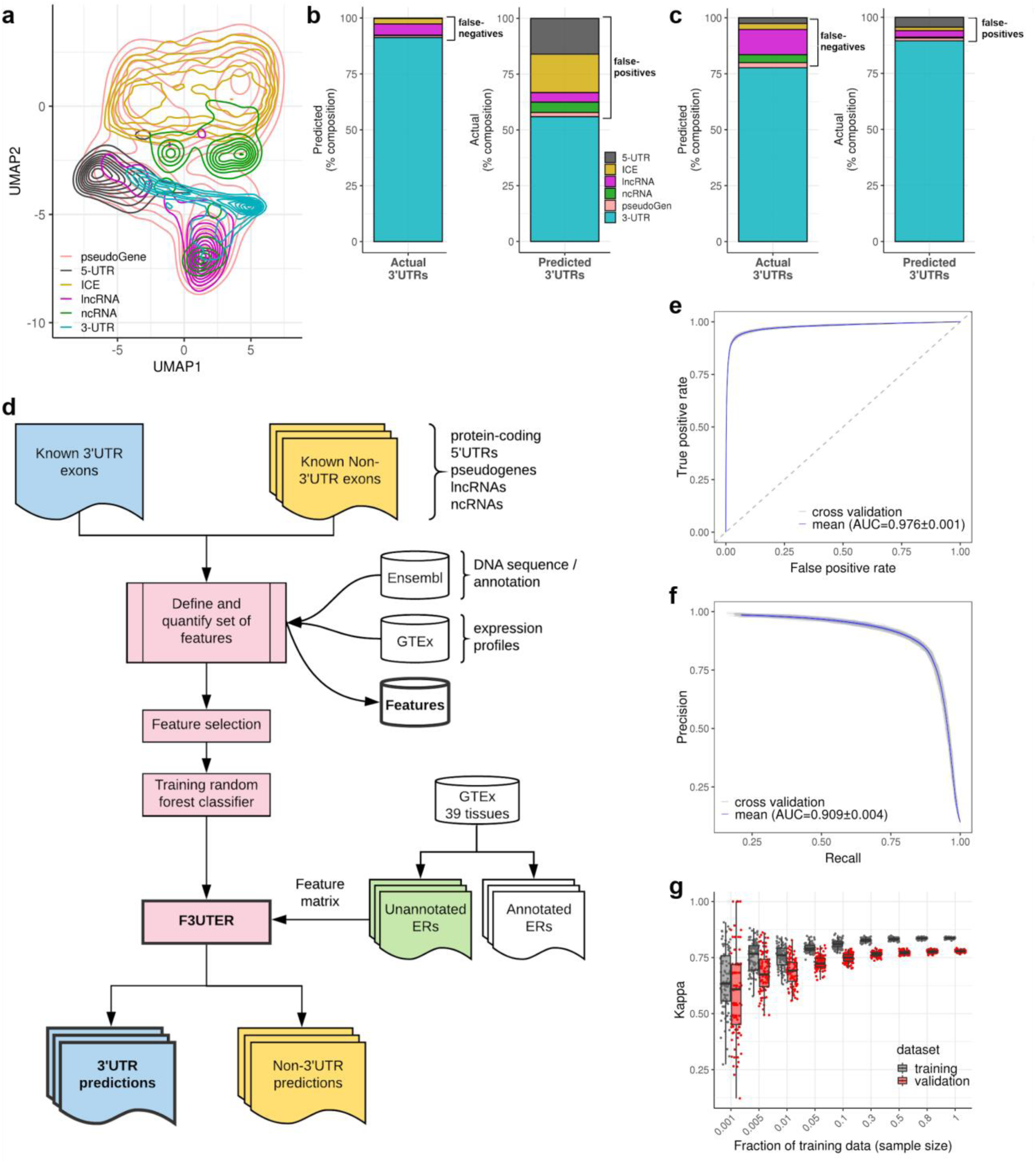
Classification of 3’UTRs from other transcribed elements in the genome. **(a)** UMAP representation of features, with elements labelled by genomic classes. **(b)** Classification of 3’UTRs using an elastic net multinomial logistic regression. **(c)** Classification of 3’UTRs using a multinomial random forest classifier. **(d)** General framework of F3UTER: the core of the framework is a random forest classifier trained on omic features derived from known 3’UTRs and non-3’UTRs. The omic features are based on either genomic (DNA sequence) or transcriptomic (RNA-seq from GTEx) properties. To make predictions, genomic coordinates of ERs are given as input, from which a feature matrix is constructed. The output of the framework is ERs categorised into potential 3’UTRs and non-3’UTRs with their associated prediction probability scores. **(e, f)** ROC and precision recall curves of F3UTER evaluated using 5-fold cross validation. **(g)** Bias- variance trade-off plot demonstrating the performance of F3UTER on training and validation datasets grouped by the sample size of the training data.

### F3UTER accurately distinguishes 3’UTRs from other genomic elements

Next, we measured the predictive value of the omic features we had identified to distinguish between unannotated 3’UTRs and other expressed elements if used collectively. We trained an elastic net multinomial logistic regression model and evaluated its performance using 5-fold cross validation repeated 20 times (**Methods**). Taking all classes into account, the multinomial logistic regression model achieved an accuracy of 74% and a kappa of 0.52 in distinguishing between the different genomic classes. Consistent with the UMAP visualisation, we found that known 3’UTRs were most likely to be misclassified as lncRNAs (4.98%), followed by ICEs (2.46%) and pseudogenes (0.88%) (**Figure 2b**). On the other hand, false-positive 3’UTR predictions, which totalled 44%, were predominantly composed of known ICEs (17.23%) and 5’UTRs (16.06%) (**Figure 2b**).

Since the high false-positive rate of our multinomial logistic regression model would be a significant barrier to reliably predict unannotated 3’UTRs from intergenic ERs, we generated an alternative machine-learning-based approach to address this problem. The resulting random forest multinomial classifier was assessed for its performance using 5-fold cross validation repeated 20 times (**Methods**). We found that the random forest multinomial classifier had a significantly higher accuracy (76%; *p* < 2.2 × 10^−16^, Wilcoxon Rank Sum Test) and kappa (0.56; *p* < 2.2 × 10^−16^, Wilcoxon Rank Sum Test) in comparison to the multinomial logistic regression model (**Supplementary Figure S3**). While the false-negative rate was higher (random forest classifier rate of 22%; logistic regression rate of 9%, **Figure 2c**), importantly the random forest-based classifier reduced false-positive calling of 3’UTRs to 10% (4.4% 5’UTR, 2.7% lncRNA, 1.5% ICE and 1.2% pseudogenes) compared to 44% using logistic regression. We also simplified the classification problem to a binary one and generated a second random forest classifier, aiming only to distinguish between 3’UTRs and non-3’UTRs. This resulted in the development of our final random forest classifier, **F**inding **3**’ **U**n-**t**ranslated **E**xpressed **R**egions (F3UTER, **Figure 2d**).

To assess F3UTER’s performance, we performed 5-fold cross validation (repeated 20 times) and calculated metrics such as accuracy, sensitivity, specificity, kappa, area under the ROC curve (AUC-ROC) and area under the precision-recall curve (AUC-PR). F3UTER achieved a mean accuracy of 0.96, sensitivity of 0.92, specificity of 0.96, kappa of 0.78, AUC-ROC of 0.98 (**Figure 2e**) and AUC-PR of 0.91 (**Figure 2f**) on the validation datasets (hold out). We found that F3UTER performed similarly on both the training and validation datasets in the cross validation (**Figure 2g**). In addition, increasing the sample size of training data reduced the variability in model predictions and hence, made it more stable. Taken together, these findings suggested that we were not overfitting the classifier. Finally, we investigated the contributions of individual features towards the accuracy and node homogeneity (Gini coefficient, **Methods**) of 3’UTR classification. Interestingly, we found that features derived directly from sequence data (e.g. conservation and PAS) as well as from the transcriptomic data, namely mean-PD and mean-EE (**Supplementary Figure S4**), most significantly contributed to the accuracy of F3UTER. This shows that F3UTER leverages both genomic and transcriptomic features to classify 3’UTRs, which would be expected to enable the identification of tissue-specific unannotated 3’UTRs.

### Evaluation of F3UTER using 3’-end sequencing data validates unannotated 3’UTR predictions

We evaluated the performance of F3UTER using an independent dataset consisting of both RNA-seq data and paired 3’-seq in B cells [23]. The latter, a form of 3’-end sequencing, was performed to identify poly(A) sites experimentally. Since poly(A) sites are present at the very end of 3’UTRs, unannotated 3’UTRs should overlap or be in the close vicinity of a poly(A) site. It should be noted that unlike the GTEx RNA-seq dataset which we used for our previous analyses and which consists of hundreds of samples for most tissues, this B cell dataset consisted of only two RNA-seq samples. Since detecting unannotated ERs relies on averaging RNA-seq coverage across many samples to reduce the contribution of transcriptional noise to ER definition, calling ERs from only two samples would likely result in inaccuracies at ER boundaries. Although this would be expected to significantly reduce the confidence in the detection of unannotated ERs and potentially underestimate the performance of F3UTER, the paired RNA-seq and 3’-seq nature of this B cell dataset enabled us to confidently validate 3’UTR predictions using gold standard experimental data.

First, we identified 3’ unannotated intergenic ERs in B cells from the RNA-seq data following the pipeline used by Zhang et al. [15]. Then we used F3UTER to predict unannotated 3’UTRs in this B cell ER dataset, and compared these predictions to intergenic poly(A) clusters detected using 3’-seq (**Figure 3a**). We focused on confident 3’UTR predictions, defined as those with a prediction probability of > 0.6. ERs predicted to be 3’UTRs which also overlapped with a poly(A) cluster were considered to be validated, as exemplified by the intergenic ER predicted to be a novel 3’UTR of the gene *CYTIP* (**Figure 3b**). As a reference, we noted that 87.9% of known 3’UTRs overlapped with a poly(A) cluster in B cell. We found that on average, 38.5% of 3’UTR predictions were validated. This was 17.5-fold higher than that for randomly selected intergenic regions (2.2%, *p* < 0.0001, permutation test; **Supplementary Figure S5**) and 2.2-fold higher than the validation rate of non-3’UTR predictions (17.4%, **Figure 3c**). Overall, these observations demonstrate the accuracy of F3UTER and show that it can effectively distinguish unannotated 3’UTRs from other functional genomic elements in the genome.

**Figure 3.**
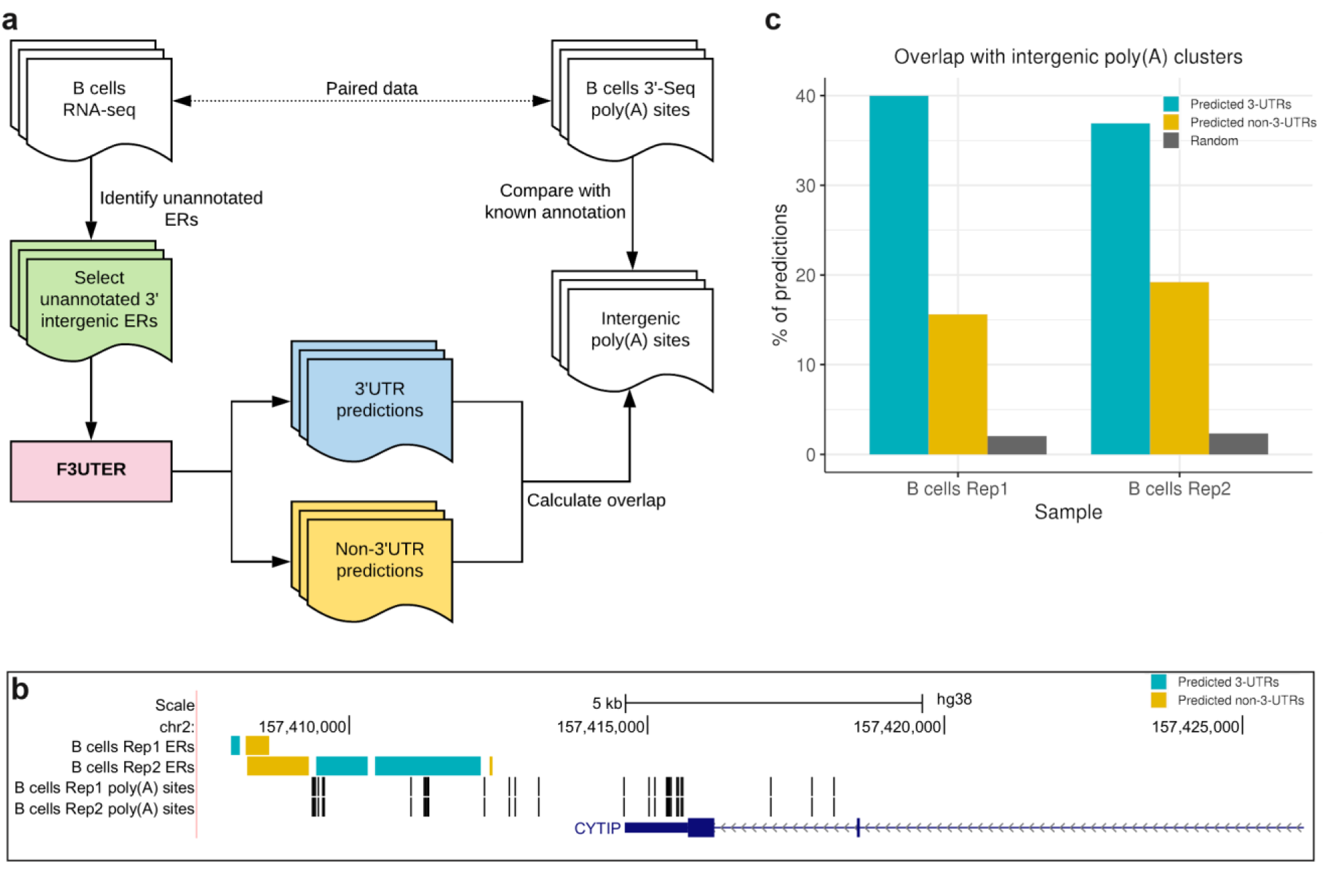
Evaluation of F3UTER predictions on an independent ER dataset. **(a)** Schematic describing the framework of the process implemented to evaluate the performance of F3UTER on ERs in B cells. **(b)** Genome browser view of the CYTIP locus, showing intergenic ERs detected downstream of CYTIP and poly(A) sites in B cells. **(c)** Bar plots showing the overlap between predictions made by F3UTER and intergenic poly(A) sites from 3‘-end sequencing in B cells. The bar for random predictions represents the mean overlap (from 10,000 permutations) between randomly selected intergenic ERs and intergenic poly(A) sites.

### Applying F3UTER across 39 GTEx tissues identifies hundreds of unannotated 3’UTRs with evidence of functional significance

We applied F3UTER to 3’ unannotated intergenic ERs identified by Zhang and colleagues [15] in 39 tissues using RNA-seq data provided by GTEx. Similar to the B cell ER dataset, we focused on confident 3’UTR predictions with a prediction probability of > 0.6 (**Supplementary File 1**). Across all tissues, we found that on average 7.9% of analysed ERs were predicted as unannotated 3’UTRs, with 8.2% being called in brain (**Supplementary Figure S6**). This equated to an average of 187 potentially unannotated 3’UTRs per tissue (ranging from 96 in adipose-subcutaneous to 348 in hypothalamus, **Figure 4a**), covering 58 to 265 kb of genomic space (mean across tissues = 138 kb, **Figure 4b**). By assigning predicted 3’UTRs to protein-coding genes either through the existence of junction reads or by proximity, we estimated that 1,513 distinct genes in total had unannotated 3’UTRs with an average of 167 genes per tissue (**Figure 4c**). As expected, the number of predicted unannotated 3’UTRs was significantly higher in the brain relative to non-brain tissues (median values of 295 and 142 in brain and non-brain tissues respectively; *p* = 1.65 × 10^−6^, Wilcoxon Rank Sum Test). This was associated with a significantly higher total genomic space (median values of 232 kb and 104 kb in brain and non-brain tissues respectively; *p =* 1.43 × 10^−8^, Wilcoxon Rank Sum Test) and higher number of implicated genes (median values of 270 and 127 in brain and non-brain tissues respectively; *p =* 1.65 × 10^−6^, Wilcoxon Rank Sum Test). This data suggests that incomplete annotation of 3’UTRs is present in all human tissues but is most prevalent in the brain.

**Figure 4.**
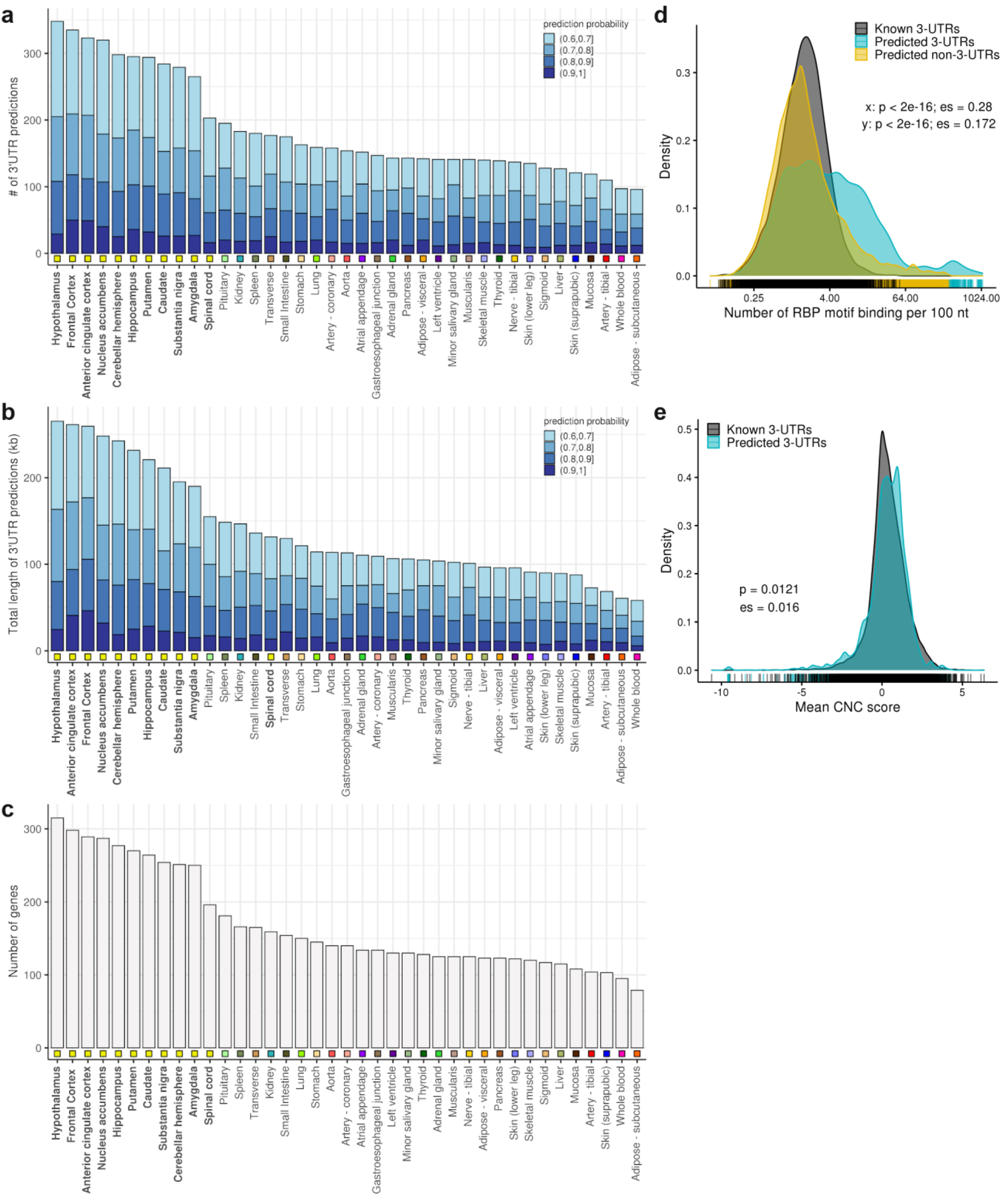
Unannotated 3’UTR predictions across 39 GTEx tissues. **(a)** Number of unannotated 3’UTRs predicted by F3UTER. **(b)** Total genomic space of unannotated 3’UTRs. **(c)** Number of genes associated with unannotated 3’UTRs. In each bar plot, tissues are sorted in descending order of the values plotted on y-axis. The square boxes below the bars are color-coded to group the tissues according to their physiology. The predictions are grouped and color-coded based on their prediction probability scores from F3UTER. **(d)** Density distributions comparing the RBP binding density across known 3’UTRs, predicted 3’UTRs and predicted non-3’UTRs. p: p-value of comparison calculated using Wilcoxon Rank Sum Test; es: effect size; x: predicted 3’UTRs vs. known 3’UTRs; y: predicted 3’UTRs vs. predicted non-3’UTRs. **(e)** Density distributions comparing the “constrained non-conserved” (CNC) scores between known and predicted 3’UTRs. p: p-value of comparison calculated using Wilcoxon Rank Sum Test; es: effect size.

Next, we investigated the functional significance of unannotated 3’UTRs by analysing their potential interaction with RBPs. This in silico analysis was performed because selective RBP binding at 3’UTRs is thought to be key in explaining the selection of alternate PASs and its impact on mRNA stability and localisation [24]. Using the catalogue of known RNA binding motifs from the ATtRACT database [25], we examined the binding density of 84 RBPs across all unannotated 3’UTRs (**Methods**). Consistent with previous reports demonstrating higher RBP binding densities in known 3’UTRs relative to other genomic regions [26], we found that 3’UTR predictions were enriched for RBP binding motifs compared to non-3’UTR predictions (*p* < 2.2 × 10^−16^, effect size (es) = 0.17, Wilcoxon Rank Sum Test, **Figure 4d**). Surprisingly, we noted that unannotated 3’UTRs were also enriched for RBP binding motifs compared to known 3’UTRs (*p* < 2.2 × 10^−16^, es = 0.28, Wilcoxon Rank Sum Test, **Figure 4d**) suggesting that these regions may be of particular functional significance. To investigate this further, we leveraged constrained, non-conserved (CNC) scores [27], a measure of human-lineage-specificity, to determine whether the unannotated 3’UTRs identified were of specific importance in humans. CNC score, a metric combining cross-species conservation and genetic constraint in humans, was used to identify and score genomic regions which are amongst the 12.5% most constrained within humans but yet are not conserved. We found that unannotated 3’UTRs exhibited higher CNC scores compared to known 3’UTRs (*p =* 0.012, es = 0.016, Wilcoxon Rank Sum Test, **Figure 4e**). Thus, together our analyses suggested that unannotated 3’UTRs are not only functionally important but may be particularly crucial in human-specific biological processes.

### F3UTER identifies unannotated 3’UTRs of genes associated with synaptic function

Given the evidence for the functional importance of unannotated 3’UTRs predicted by F3UTER, we wanted to explore their biological relevance. To do this, we began by categorising all unannotated 3’UTRs into four sets based on their tissue-specificity: absolute tissue-specific, highly brain-specific, shared and ambiguous (**Methods** and **Supplementary Figure S7**). Using this non-redundant set of 3’UTRs, we found that on average, we extended the current annotation per gene by 681 bp in highly brain-specific (1.4x the known maximal 3’UTR length), 633.6 bp in tissue-specific (0.95x the known maximal 3’UTR length), and 496.63 bp in shared predictions (0.88x the known maximal 3’UTR length) respectively. Next, we repeated the RBP and CNC analysis for each category finding that all unannotated 3’UTR sets showed significant enrichment of RBP binding motifs when compared not only to non-3’UTR predictions (*p* ≤ 2.5 × 10^−5^, Wilcoxon Rank Sum Test), but also to known 3’UTRs (*p* ≤ 3.9 × 10^−7^, Wilcoxon Rank Sum Test), with the brain-specific set having the largest effect size (es ≥ 0.17) (**Figure 5a**). Focussing on CNC scores, we found that while none of the unannotated 3’UTR sets showed significant differences in score compared to known 3’UTRs (*p* ≥ 0.121, Wilcoxon Rank Sum Test), brain-specific unannotated 3’UTRs trended to significance with the largest effect size relative to other sets (**Figure 5a**). Together, these observations lead us to conclude that highly brain-specific 3’UTR predictions were likely to be of most biological interest.

**Figure 5.**
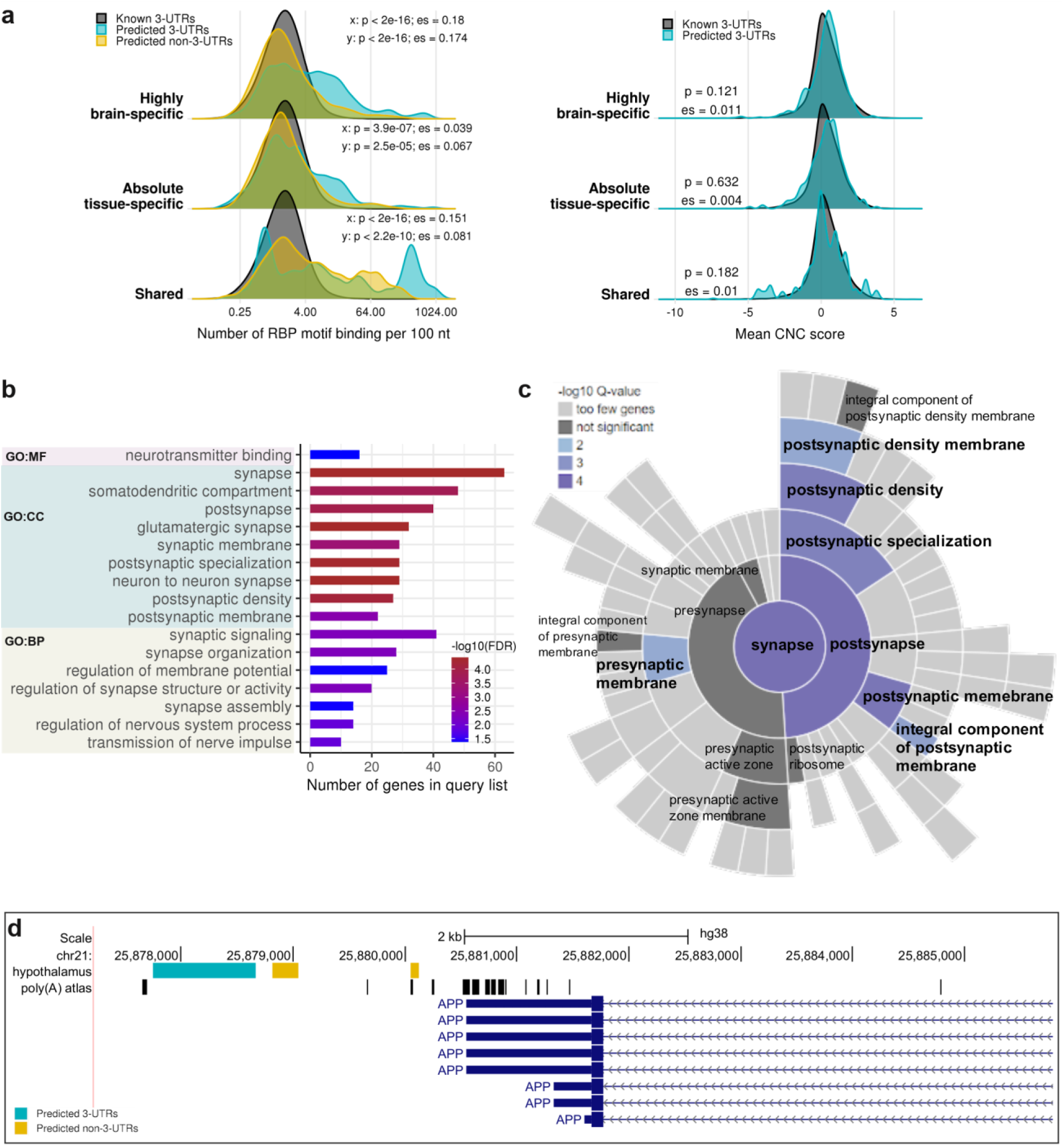
Functional significance of highly brain-specific unannotated 3’UTRs. **(a)** Density distributions comparing RBP binding and “constrained non-conserved” (CNC) scores between known, predicted 3’UTRs and predicted non-3’UTRs, categorised according to their tissue- specificity. p: p-value of comparison calculated using Wilcoxon Rank Sum Test; es: effect size; x: predicted 3’UTRs vs. known 3’UTRs; y: predicted 3’UTRs vs. predicted non-3’UTRs. **(b)** GO terms enriched amongst the list of genes associated with highly brain-specific unannotated 3’UTRs. MF: molecular function; CC: cellular component; BP: biological process. **(c)** Sunburst plot showing the cellular component SynGO terms over-represented in genes associated with highly brain-specific 3’UTRs. The inner rings of the plot represent parent terms, while outer rings represent their more specific child terms. Rings are colour coded based on the enrichment q-value of the terms. **(d)** Genome browser view of the APP locus, showing intergenic ERs detected downstream of APP in the hypothalamus, and poly(A) sites from the poly(A) atlas data.

These observations raised the question of what types of genes are associated with highly brain-specific 3’UTR predictions. Interestingly, we found that while genes linked to brain-specific non- 3’UTR predictions had no GO term enrichments, those linked to an unannotated brain-specific 3’UTR were significantly enriched for synaptic function (“synaptic signalling”, “synapse organisation” and “protein localization to postsynaptic specialization membrane”; *q* = 4.97 × 10^−3^) (**Figure 5b, Supplementary File 2**). Using SynGO (the synaptic GO database [28]) to obtain more granular information, we found that genes associated with unannotated 3’UTRs were more significantly enriched for terms relating to post-synaptic (“protein localisation in postsynaptic density”, *q* = 2.87 × 10^−4^; postsynaptic function, *q* = 4.1 × 10^−3^), as compared to presynaptic structures (“localisation in presynapse”, *q* = 0.03; presynaptic function, *q* = 0.1) (**Figure 5c**, **Supplementary File 2**). Furthermore, we found that genes linked to unannotated brain-specific 3’UTRs were significantly enriched for those already associated with rare neurogenetic disorders (*p =* 0.01, hypergeometric test) and more specifically adult-onset neurodegenerative disorders (*p =* 0.03, hypergeometric test). For example, we detected an unannotated 3’UTR in the brain linked to the gene, *APP*, a membrane protein which when mutated gives rise to autosomal dominant Alzheimer’s disease and encodes for amyloid precursor protein, the main constituent of amyloid plaques [29]. We detected a 920 bp long brain-specific unannotated 3’UTR located 1.8 kb downstream of *APP* (**Figure 5d**) and only 51 bp from an intergenic poly(A) site on the same strand as *APP* gene as reported by the poly(A) atlas. Other similar examples included the genes, *C19orf12*, *RTN2*, *SCN2A* and *OPA1* (**Supplementary Figures S8 & S9**).

### Brain-specific unannotated 3’UTRs interact with RBPs implicated in neurological disorders

Next, we investigated the information content of brain-specific unannotated 3’UTRs by comparing RBP binding enrichments between brain-specific and shared 3’UTR predictions (**Methods**). By using shared 3’UTR predictions as the negative control, we removed RBPs associated with non-brain tissues and so identified RBP binding of greatest relevance to human brain function. This analysis identified 22 RBPs with significantly enriched binding in the brain-specific unannotated 3’UTRs (*adjusted p* < 10^−5^) (**Supplementary Table 1**). We found that nine of these RBPs were previously known to be associated with “mRNA 3’UTR binding” (*q* = 2.23 × 10^−14^, **Supplementary File 3**), including *TARDBP,* an RNA binding protein implicated in both frontotemporal dementia and amyotrophic lateral sclerosis [30]. Of the 75 gene targets that we identified for *TARDBP* through unannotated 3’UTRs, up to 50 were known to be *TARDBP* targets based on computational scanning of existing 3’UTR annotations for *TARDBP* motif (47%, *p =* 0.008, hypergeometric test) and iCLIP experiments (44%, *p =* 1.47 × 10^−6^, hypergeometric test). However, this implied that 25 gene targets were not previously known to harbour *TARDBP* binding motifs based on current annotation. Another RBP which was identified to be significantly enriched in brain-specific unannotated 3’UTRs was *RBFOX1* (*adjusted p* = 1.78 × 10^−18^), a neuronal splicing factor implicated in the regulation of synaptic transmission [31] and whose mRNA targets have been implicated in autism spectrum disorders [32]. We identified 89 gene targets with a predicted *RBFOX1* binding motif within their associated unannotated 3’UTRs. Of these 89 genes, only 31 (35%) had a predicted *RBFOX1* binding motif within their existing 3’UTRs, again implying that unannotated 3’UTRs provide valuable novel binding sites. Furthermore, GO and SynGO enrichment analyses (**Supplementary File 3**) demonstrated that the target genes of *RBFOX1* were significantly enriched for synaptic (“synaptic membrane adhesion”, *q* = 1.58 × 10^−2^) and postsynaptic terms (“postsynapse”, *q* = 0.01), consistent with the previously known functions of *RBFOX1* [31]. These results show that the identification of brain-specific unannotated 3’UTRs can recognise new genes within known regulatory networks, which can provide novel, disease-relevant insights.

## Discussion

In this study we generate a machine learning-based classifier, F3UTER, which leverages transcriptomic as well as genomic data to predict unannotated 3’UTRs. F3UTER outperforms a state-of-the-art statistical learning approach, elastic net logistic regression, whilst retaining its interpretability capabilities. We apply F3UTER to transcriptomic data covering 39 human tissues studied within GTEx, enabling the identification of tissue-specific unannotated 3’UTRs. Using this large, public, short-read RNA-seq data set, we predict unannotated 3’UTRs for 1,513 genes, (equating to 5.4 Mb of genomic space in total across 39 tissues) and demonstrate that F3UTER can be successfully applied to human genomic regions from any tissue with existing bulk RNA-seq data. In fact, even though intergenic ERs in B cells were generated using only two samples, we were able to validate 38.5% of the unannotated 3’UTR predictions using 3’-end sequencing data, showing that F3UTER can be a useful tool even for small RNA-seq datasets. Furthermore, it should be noted that F3UTER does not depend on ER datasets as input, but instead any set of interesting human genomic regions can be used. Given the continued popularity and high availability of short-read RNA-seq data across tissues, cell types and disease states, we believe that (1) F3UTER could be applied more broadly to improve our understanding of 3’UTR diversity and usage, and (2) the set of omic features devised within this study could form the basis for other predictive models aimed at increasing the accuracy of human transcriptomic annotation.

We focus on F3UTER-predicted 3’UTRs in human brain, which we find to be most prevalent when comparing predictions across all 39 human tissues. We believe that the higher frequency of incomplete 3’UTR annotation in human brain could be attributed to several factors including: (1) higher transcript diversity with many rare isoforms expressed in this tissue; (2) high cellular heterogeneity complicating detection of tissue- /cell-type specific transcripts; (3) historically lower availability of human brain samples; and (4) reliance on post-mortem tissues, which suffer from RNA degradation resulting in decreased accuracy of transcript identification.

While we find that collectively the unannotated 3’UTRs predicted by F3UTER were significantly enriched for RBP binding and exhibited high human lineage-specificity, the latter was primarily driven by brain-specific 3’UTR predictions. Overall, these findings suggest that predicted 3’UTRs are likely to be functionally important in the human genome. Moreover, these findings provide some explanation for the difficulties of identifying 3’UTRs through cross-species analyses particularly when considering brain-specific transcripts. Interestingly, we find that brain-specific unannotated 3’UTRs were enriched for binding of RBPs already implicated in neurological disorders, such as *TARDBP* and *RBFOX1.* Furthermore, genes linked to unannotated brain-specific 3’UTRs were significantly enriched for those already associated with rare neurogenetic and adult-onset neurodegenerative disorders, and for genes involved in synaptic function.

Taken together, our results demonstrate that F3UTER not only improved 3’UTR annotation, but also identified unannotated 3’UTRs in the human brain which provided novel insights into the mRNA-RBP interactome with implications for our understanding of neurological and neurodevelopmental diseases. With this in mind, we note the growing interest in the role of 3’UTR-based mechanisms in translational regulation within complex, large, polarised cell types such as neurons [4, 5, 33, 34]. Although increasing use of single-nuclei RNA-seq, together with long-read RNA-seq will provide further insights into alternative 3’UTR usage and will impact the field considerably, these technologies still have significant limitations for the identification of rare transcripts. Therefore, we believe that F3UTER, which can effectively utilise existing short-read RNA-seq data sets, will be of interest to a wide range of researchers. Furthermore, we release our results through an online resource (F3UTER: https://astx.shinyapps.io/F3UTER/) which allows users to both easily query unannotated 3’UTRs and inspect the omic features driving the classifier’s prediction for an ER of interest.

## Methods

### ER data

We collected the set of intergenic ERs identified by Zhang and colleagues [15] in 39 GTEx tissues, comprising of 11 non-redundant brain tissues and 28 non-brain tissues (total intergenic ERs = 9,339,770). Each ER was associated to a protein-coding gene by extracting genes which connected to the ER via a junction read. In cases where no junction read was present, the nearest protein-coding gene was assigned to the ER. From this dataset, we selected intergenic ERs located within 10 kb of their associated gene, resulting in 237,540 ERs. In this dataset, 4% of the ERs were associated to a gene via a junction read. Based on the location of intergenic ERs with respect to their associated genes, i.e. whether upstream or downstream, we annotated their orientation as 5’ (92,148 ERs) or 3’ (145,392 ERs) respectively. The total genomic space of these intergenic ERs was calculated by adding the length of all ERs in each tissue. To further remove ERs which were unlikely to be 3’UTRs, we selected 3’ intergenic ERs with a length ≤ 2 kb – which is the third quartile limit of known 3’UTR exon lengths. We also removed small ERs with length ≤ 40 nucleotides (nt) for which feature calculation can be problematic. This resulted in a set of 93,934 ERs across all 39 tissues, and this set was used as input to F3UTER.

### Assembling positive and negative 3’UTR learning datasets

For positive examples, we used known 3’UTRs, while for negative examples, we used regions in the genome which are known to be non-3’UTRs, namely 5’UTRs, internal coding exons (ICEs), lncRNAs, ncRNAs and pseudogenes. Ensembl human genome annotation (v94 GTF) was used to extract the genomic coordinates of these different genomic classes. For all classes in our training dataset, firstly, we selected high confidence annotations at the transcript level with transcript support level (TSL) = 1. Secondly, we collapsed and combined multiple transcripts associated with a single gene to make a consensus “meta-transcript” per gene. This merged all the overlapping regions emerging from the same gene. Finally, we extracted exons with width >= 40 (nt) from these meta-transcripts to serve as learning examples.

To capture regions of 3’UTR exons, 5’UTR exons and ICEs, transcripts from protein-coding genes were selected. For ICE examples, transcripts with at least three coding exons were further selected (as transcripts with less than three exons would not contain an internal exon) and their first and last coding exons were removed to capture ICEs. To capture lncRNA, ncRNA and pseudogene exons, we selected annotations from the GTF file with the following gene biotypes:

- **lncRNA:** “non_coding”, “3prime_overlapping_ncRNA”, “antisense”, “lincRNA”, “sense_intronic”, “sense_overlapping”, “macro_lncRNA”
- **ncRNA:** “miRNA”, “misc_RNA”, “rRNA”, “snRNA”, “snoRNA”, “vaultRNA”
- **pseudogene:** “pseudogene”, “processed_pseudogene”, “unprocessed_pseudogene”, “transcribed_processed_pseudogene”, “transcribed_unitary_pseudogene”, “transcribed_unprocessed_pseudogene”, “translated_processed_pseudogene”, “unitary_pseudogene”, “unprocessed_pseudogene”, “TR_V_pseudogene”, “TR_J_pseudogene”, “rRNA_pseudogene”, “polymorphic_pseudogene”, “IG_V_pseudogene”, “IG_pseudogene”, “IG_J_pseudogene”, “IG_C_pseudogene”

### Calculating omic features

For each region in the training dataset, we calculated several genomic and transcriptomic based features. Transcriptomic features were used to account for tissue-specific properties of transcribed elements in the genome.

#### Genomic (sequence) based features

● **Poly(A) signals (number of features, n=1):** Previous studies have shown that 3’UTR sequences of most mammalian genes contain the consensus AAUAAA motif (or a close variant) 10-30 nt upstream of the poly(A) site [8]. These motif sites are recognised and bound by the cleavage and polyadenylation specificity factor (CPSF), and are referred to as polyadenylation signals (PASs). PASs are an important characteristic of 3’UTRs and are involved in the regulation of the polyadenylation process [8]. We used 12 commonly occurring PASs (AAUAAA, AUUAAA, AGUAAA, UAUAAA, AAUAUA, AAUACA, CAUAAA, GAUAAA, ACUAAA, AAUAGA, AAUGAA, AAGAAA) [9, 12, 35, 36] to construct a consensus position weight matrix (PWM). Each region was scanned for potential PWM matches and a binary outcome was reported i.e. whether the region contains a potential PAS or not. The “searchSeq’’ function (with min.score= “95%”) from the R package “TFBSTools” [37] was used to detect PWM matches.
● **Mono- and di-nucleotide frequency (n=20):** The sequence composition in 3’UTRs, especially near the poly(A) sites has been shown to be important for polyadenylation [8, 9, 35]. The frequency probability of each mono-nucleotide (i.e. A, T G, C; n=4) and di- nucleotide pair (n=16; e.g. AA, AT, GC, GG) was calculated as the number of nucleotide occurrences divided by the length of the region.
● **DNA sequence conservation (n=1):** Sequences of non-protein coding transcripts and un-translated regions are poorly conserved compared to protein-coding sequences [38, 39]. For every genomic position, we extracted the phastCons score of the human genome (hg38) across 7 species pre-computed by the UCSC genome browser, and averaged it across the region to calculate mean sequence conservation score for each region.
● **Transposons (n=1):** Previous studies have revealed that transposons are highly enriched within lncRNAs compared to protein-coding genes and other non-coding elements [40, 41]. These transposable elements are considered to be the functional domains of lncRNAs. We calculated the total fraction of region covered with transposons – LINEs, SINEs, LTRs, DNA and RC transposons. The hg38 genomic coordinates of the transposable elements (Dfam v2.0) were downloaded from http://www.repeatmasker.org/species/hg.html.
● **DNA structural properties (n=16):** The underlying sequence composition of a DNA molecule plays an important role in determining its structure. As a result, similar DNA sequences have a tendency to have similar DNA structures [42]. We calculated 16 properties of DNA structures which can be predicted from a nucleotide sequence based on previous experiments. To quantitatively measure a structural property from a nucleotide sequence, we used pre-compiled conversion tables downloaded from http://bioinformatics.psb.ugent.be/webtools/ep3/?conversion [43]. Depending on the structural property, we extracted scores for each di-nucleotide or tri-nucleotide occurrence in the sequence from the conversion tables, and averaged the scores across the region.

#### Transcriptomic based features

● **Entropy efficiency (n=1):** We measured the uniformity of read coverage across a region using entropy efficiency, as described in Gruber et al. [44]. The entropy efficiency (EE) of a region (x) was calculated as, 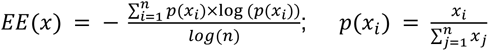 where *n*, represents the length of the region and *p*(*x_i_*) is the read count at position *i* divided by the total read count of the region. For each region, we calculated EE in 39 GTEx tissues and averaged it across all the tissues to obtain a baseline distribution of EE scores.
● **Percentage difference (n=1):** We calculated the percentage difference (PD) between the read counts at the boundaries of a region. For read counts *r*_1_ and *r*_2_ measured at the boundaries of a region *x*, PD was calculated as: 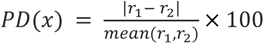. For each region, we calculated PD in 39 GTEx tissues and averaged it across all the tissues to obtain a baseline distribution of PD scores.

### Univariate and multivariate analysis

For univariate analysis, we performed non-parametric Kruskal–Wallis test and proportion Z-test for continuous and categorical variables, respectively, to identify features with significant differences across all the genomic classes. We used UMAP [22] to visualise all the features in two-dimensional space. The UMAP analysis was performed using the R package “umap” with default parameters. The clusters were visualised as a 2D density and a scatter plot. Each data point was labelled and coloured according to its genomic class.

To perform multivariate analysis, a feature matrix was generated where rows represented regions from the training dataset (n=179,968), and columns represented the quantified features (n=41). The features were scaled and centred in R using the preProcess function of R “Caret” package [45]. The elastic net multinomial logistic regression model was trained using the “glmnet” R package [46] with the following parameters: family = “multinomial”, alpha=0.5, nlambda=25 and maxit=10,000. The random forest multinomial classifier was trained within Caret using the “randomForest” package [47] with default parameters (ntree = 500, nodesize = 1). We performed a 5-fold cross validation (repeated 20 times) to evaluate the performance of these multinomial classifiers, where the model was trained on 80% of the data (training dataset) and tested on 20% of the remaining data (validation dataset). Downsampling of the data was employed to correct for imbalance in the sample size of the classes. For each cross validation run, we produced a confusion matrix for each prediction class using the Caret’s confusionMatrix function and computed the false- positive and negative rates. Additionally, we calculated Cohen’s kappa, which reports the accuracy of a model compared to the expected accuracy and is a much accurate measure of performance for imbalanced datasets. These metrics were averaged across all the cross validation runs for reporting purposes.

### F3UTER construction and evaluation

We designed F3UTER as a binary classifier to categorise an ER into a 3’UTR (positive) or a non- 3’UTR (negative). This random forest classifier was implemented in R using Caret as the machine learning framework and “randomForest” as the machine learning algorithm within Caret. The random forest classifier was trained using the default parameters (ntree = 500, nodesize = 1). We performed a 5-fold cross validation (repeated 20 times) to evaluate the performance of the F3UTER. For each cross validation run, we calculated the performance metrics such as accuracy, kappa, sensitivity, specificity, ROC curve and precision-recall curve, using the caret’s confusionMatrix function. Variable importance was measured using mean decrease in accuracy and Gini coefficient, as natively reported by random forest. The Gini coefficient measures the contribution of variables towards homogeneity of nodes in the random forest tree. These metrics were averaged across all the cross validation runs for reporting purposes. For bias-variance trade-off analysis, we trained F3UTER on sequentially increasing sample size of training data (0.1%, 0.5%, 1%, 5%, 10%, 30%, 50%, 80% and 100%), hence sequentially increasing the complexity of the model. For each sample size value, a fraction of the training data was randomly selected, and a 5-fold cross validation was performed which captured all the performance metrics for both the training and validation datasets. This process was repeated 20 times for each sample size. To make 3’UTR predictions on ER datasets, the classifier with the highest kappa statistic was selected from the cross validation process.

### Validation of 3’UTR predictions in B cells

Previously published RNA-seq and its corresponding 3’-end seq data in B cells [23] (two replicates each) was used for validating 3’UTR predictions (GEO repository: GSE111310; samples: GSM3028281, GSM3028282, GSM3028302 and GSM3028304). We processed each RNA-seq replicate individually and detected 3’ intergenic ERs using the pipeline detailed in Zhang et al. [15]. Analysed poly(A) site clusters associated with these RNA-seq samples were downloaded from poly(A) atlas [13]. These poly(A) site clusters were compared to Ensembl human genome annotation (v92) to identify sites which occur within the intergenic regions. F3UTER was applied to 3’ intergenic ERs in B cells and the resulting predictions (with prediction probability > 0.6) were compared to intergenic poly(A) site clusters to calculate their overlap. Predictions with at least a 1 bp overlap with a poly(A) site were considered to be overlapping. A permutation test was performed to inspect if the observed overlap between 3’UTR predictions and intergenic poly(A) sites is more than what we would expect by random chance. Using BEDTOOLS [48], the locations of 3’UTR predictions were shuffled in the intergenic genomic space on the same chromosome, hence generating random intergenic ERs with length, size and chromosome distribution similar to 3’UTR predictions in B cells. To shuffle the locations within the intergenic space, we excluded the genomic space covered by genes (all Ensembl bio-types) and intergenic ERs in B cells (both 3’ and 5’). The overlap between these randomly generated intergenic ERs and poly(A) sites was then calculated, and this process was repeated 10,000 times to produce a distribution of expected overlap. The p-value was calculated as 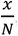, where *x* is the number of expected overlap greater than the observed overlap, and *N* is the total number of permutations. The z-score was calculated as 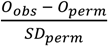 where *O_obs_* represents the observed overlap, *O_perm_* is the median of the permuted overlap, and *SD*_perm_ is the standard deviation of the permuted distribution.

### 3’UTR predictions in 39 GTEx tissues

A feature matrix of 3’ intergenic ERs was generated in each tissue. F3UTER was applied to each matrix to categorise intergenic ERs into 3’UTR (*prediction probability* > 0.60) and non-3’UTR (*prediction probability* ≤ 0.60) predictions. For each tissue, the lengths of the 3’UTR predictions were added to calculate their total genomic space (in kb). To compare brain and non-brain tissues, a two-sided Wilcoxon Rank Sum Test was applied to statistically compare the associated numbers between the two groups. To explore the biological relevance of 3’UTR predictions, they were categorised into four groups based on their tissue-specificity: absolute tissue-specific, highly brain-specific, shared and ambiguous. To do such categorisation, the genomic coordinates of ER predictions were compared across the 39 tissues. An ER which did not overlap any other ER across the tissues was labelled as “absolute tissue-specific” or present in only one tissue. On the other hand, for an ER which overlapped (≥ 1 bp) ERs from other tissues, we calculated the proportion of brain tissues in which the ER was detected. If more than 75% of the tissues were brain related, the ER was labelled as “highly brain-specific”. From the remaining data, ERs detected in at least five tissues, with their start and end coordinates within a 10 bp window, were labelled as “shared”. All the remaining ERs which did not fall in any of the above categories were labelled as “ambiguous”.

### RBP and CNCR analysis

The position weight matrices (PWMs) of RBP binding motifs in humans were collected from the ATtRACT database [25]. Motifs with less than 7 nt in length and with a confidence score of less than one, were removed to reduce false-positives in the motif matches. To remove redundancy between multiple motifs of a RBP, we further selected the longest available motif. This resulted in 84 unique PWMs, which were then used for identifying potential RBP binding using tools from the MEME suite [49]. We used FIMO [50] with a uniform background to scan query regions for potential RBP motif matches. For each RBP motif and query sequence pair, we calculated normalised counts as the number of motif matches (with *p <* 10^−4^) per 100 nt of query sequence. To summarise this analysis, we then calculated an overall RBP binding score for each query sequence by adding the normalised counts across all the RBPs. We used AME [51] with default parameters to compare binding enrichment of RBPs between highly brain-specific (query) and shared 3’UTR predictions (control). RBP motifs with an enrichment adjuested *p* – value < 10^−5^ were considered to be significantly over-represented in highly brain-specific 3’UTR predictions compared to shared 3’UTR predictions. Previously reported gene targets of TARDBP identified using iCLIP technology were extracted from the POSTAR2 database [52].

The CNC scores, as reported by Chen et al. [27], were used to quantify the occurrence of CNCRs within unannotated 3’UTRs. We extracted the CNC score for each 10 bp window and averaged it across the query region to calculate a mean CNC score for each query region.

### Calculating gene enrichment

To investigate molecular functions and biological processes significantly associated with a gene list, we performed GO enrichment analysis using the ToppFun tool in the ToppGene suite [53]. GO terms attaining an enrichment q-value (false-discovery rate computed using Benjamini-Hochberg method) < 0.05 were considered significant. Similarly, SynGO [28] was used to identify enriched GO terms (q-value < 0.05) associated with synaptic function. To calculate enrichment of genes associated with rare neurogenetic disorders, OMIM [54] genes related to neurological disorders were used (1,948 genes). The list of genes associated with adult-onset neurodegenerative disorders was extracted from Genomic England Panel App (254 green labelled genes) [55]. A hypergeometric test was used to calculate the enrichment using the total number of protein-coding genes (22,686) as the ‘gene universe’.

## Data availability

Code used to perform analyses in this study is publicly available at https://github.com/sid-sethi/F3UTER. Accession numbers of all data used in this study are listed in methods.

## Acknowledgements

We thank Matthew Davis, Greg O’Sullivan and Carla Bento for their thoughtful feedback on this study. This work was funded by a postdoctoral fellowship awarded to S.S. under the “Sustaining Innovation Postdoctoral Training Program” at Astex Pharmaceuticals (S.S., H.S.). D.Z., S.G-R., and M.R. were supported by the award of a Tenure Track Clinician Scientist Fellowship to M.R. (MR/N008324/1). Z.C. was supported through the award of a Leonard Wolfson Doctoral Training Fellowship in Neurodegeneration. S.G. was supported through the award of an Alzheimer’s Research UK PhD fellowship. J.A.B. was supported by the Science and Technology Agency, Séneca Foundation, Comunidad Autónoma Región de Murcia, Spain, through the research project 00007/COVI/20. We thank colleagues at the University College London, University of Murcia and Astex Pharmaceuticals for helpful comments.

## Author contributions

S.S., H.S., J.A.B., M.R. conceived and designed the study. S.S. conducted all the research and data analysis. M.R., J.A.B., H.S. jointly supervised this study. D.Z., S.G. provided ER datasets in GTEx tissues and helped with the analysis of ERs. S.S. developed the F3UTER online platform. Z.C. provided help and data for the CNC analysis. S.S. wrote the manuscript with help from M.R., J.A.B., and H.S. All authors contributed, read and approved the manuscript.

## Corresponding authors

Correspondence to Juan A. Botia, Harpreet Saini and Mina Ryten.

---

**Supplementary Figure S1.**
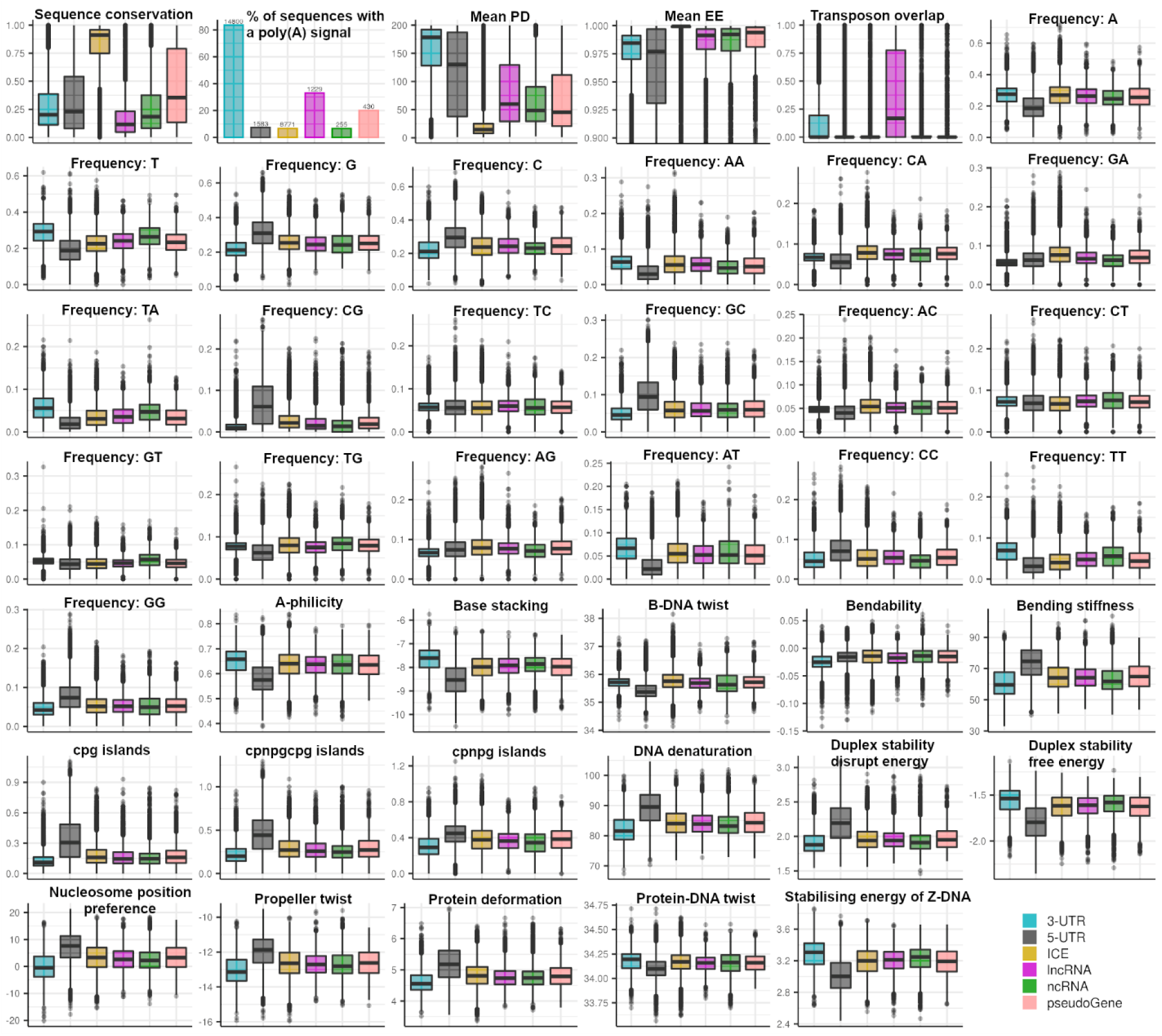
Univariate comparisons of features and genomic classes. Plots show the relationship between quantified features and genomic classes in the training dataset. A Kruskal-Wallis Test was used to compare continuous values of features across the classes, while a proportion Z-test was used for proportions. For each feature, the comparison across the classes was statistically significant with a *p* – value < 2.2 × 10^−16^.

**Supplementary Figure S2.**
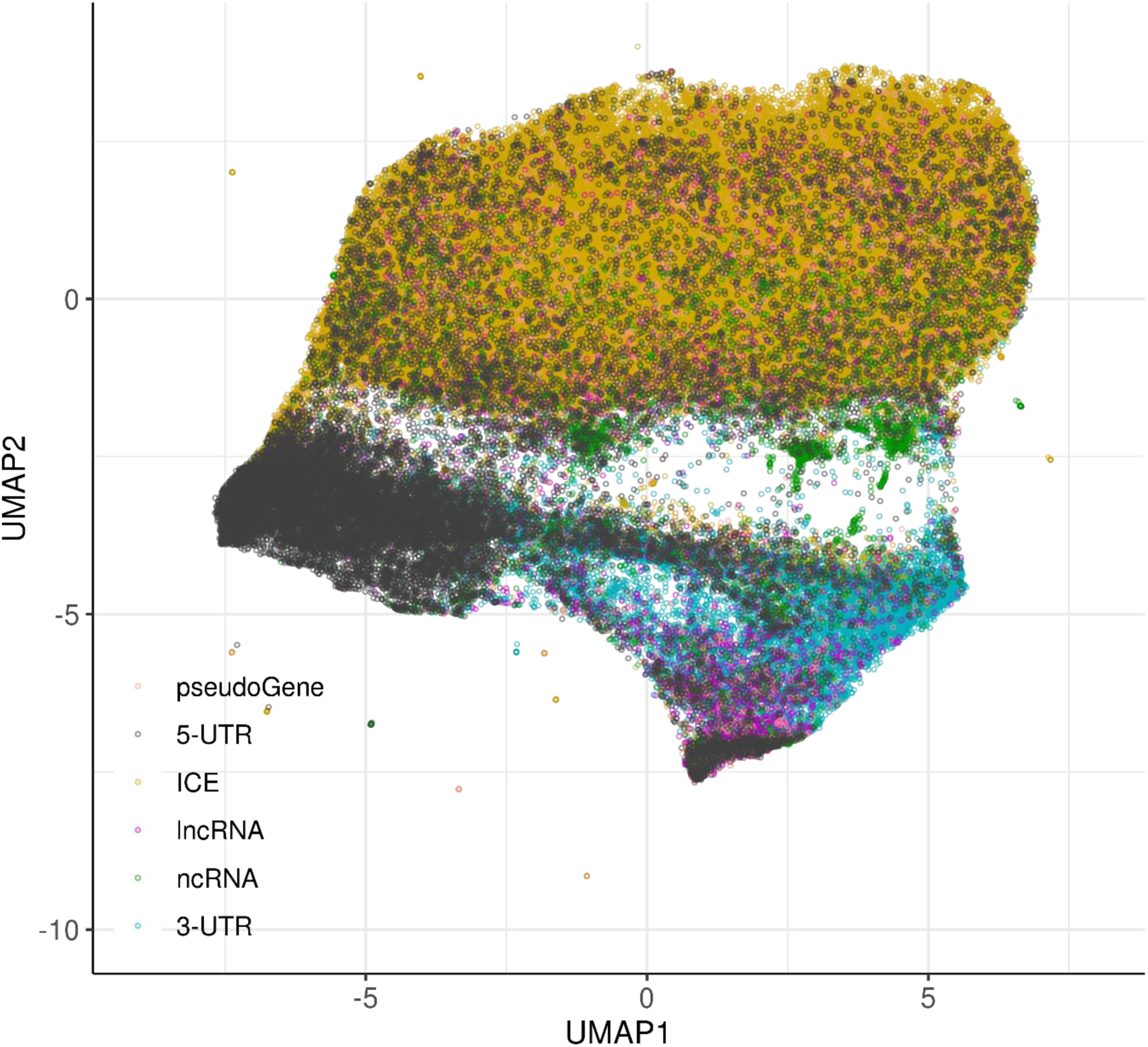
UMAP visualisation of genomic features. UMAP representation of all 41 omic features, with elements labelled by genomic classes.

**Supplementary Figure S3.**
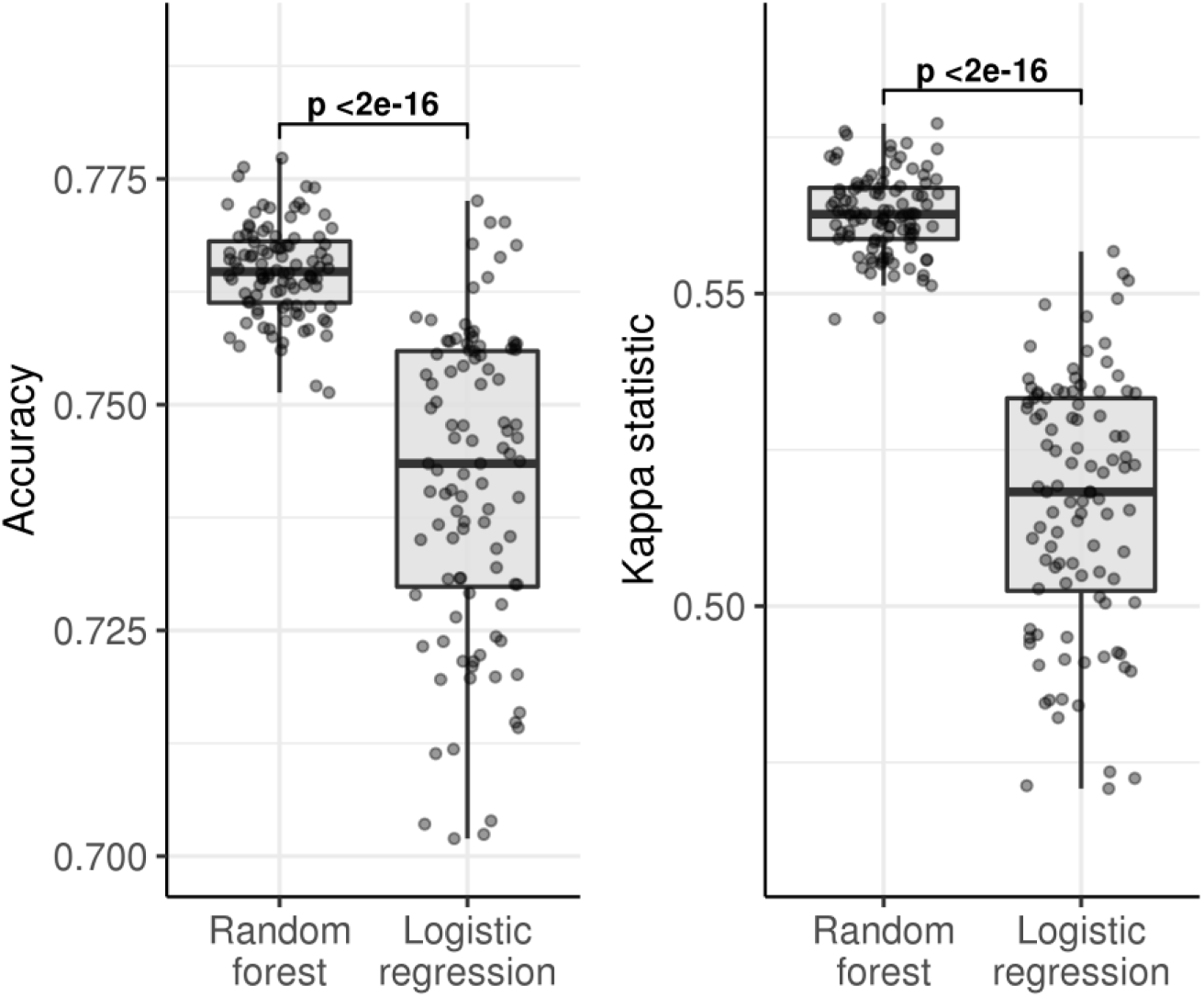
Performance of multinomial classification models measured using 5-fold cross validation repeated 20 times. Boxplots comparing the accuracy and kappa of random forest multinomial classifier and elastic net multinomial logistic regression model, to classify different genomic classes. p: p-value calculated using Wilcoxon Rank Sum Test.

**Supplementary Figure S4.**
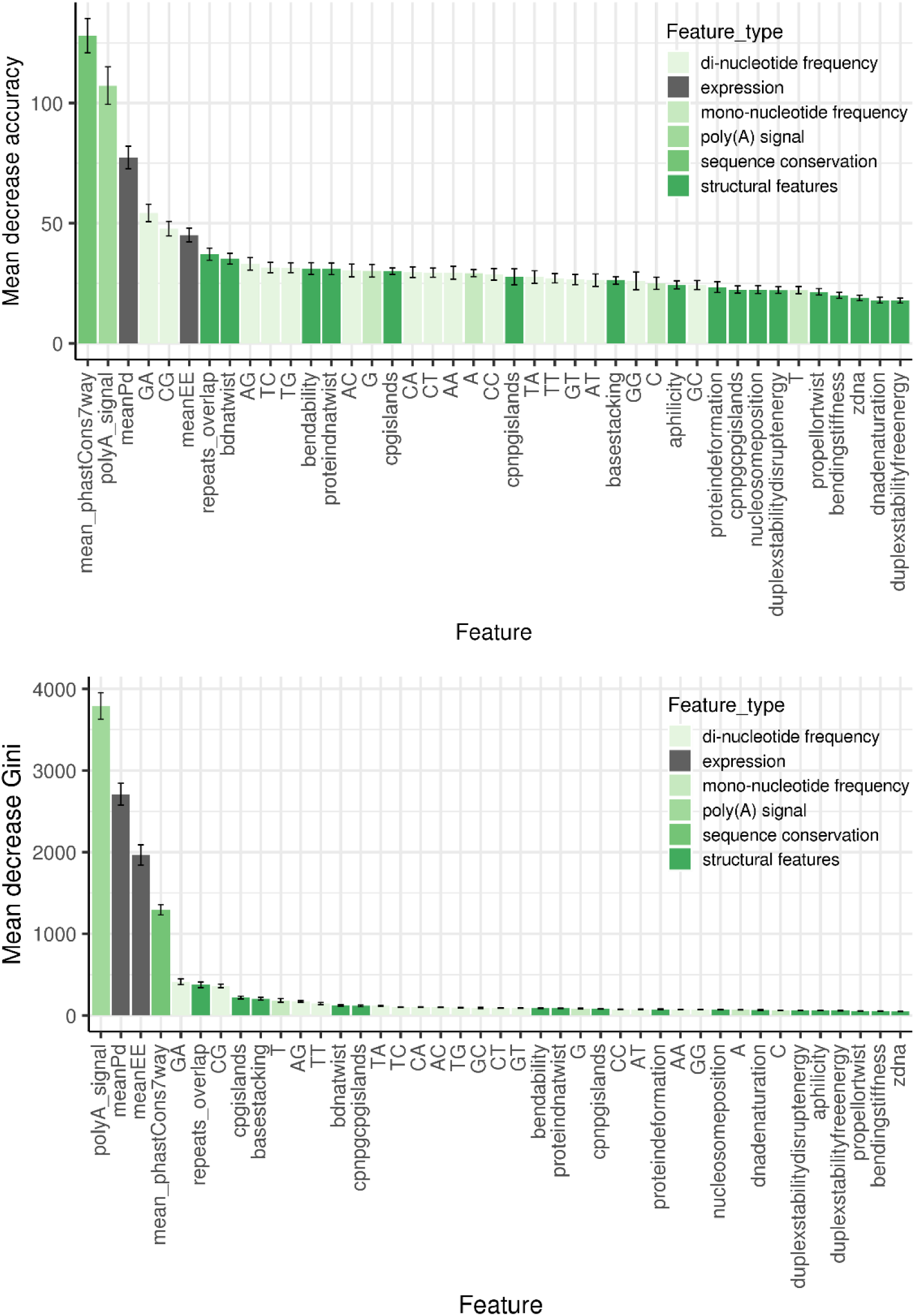
Contribution of features towards 3’UTR classification. Variable importance chart showing the importance of features in classifying 3’UTRs from other transcribed elements in the genome, as measured by mean decrease in accuracy and Gini. The features are ordered in decreasing order of their relative importance and grouped based on their type.

**Supplementary Figure S5.**
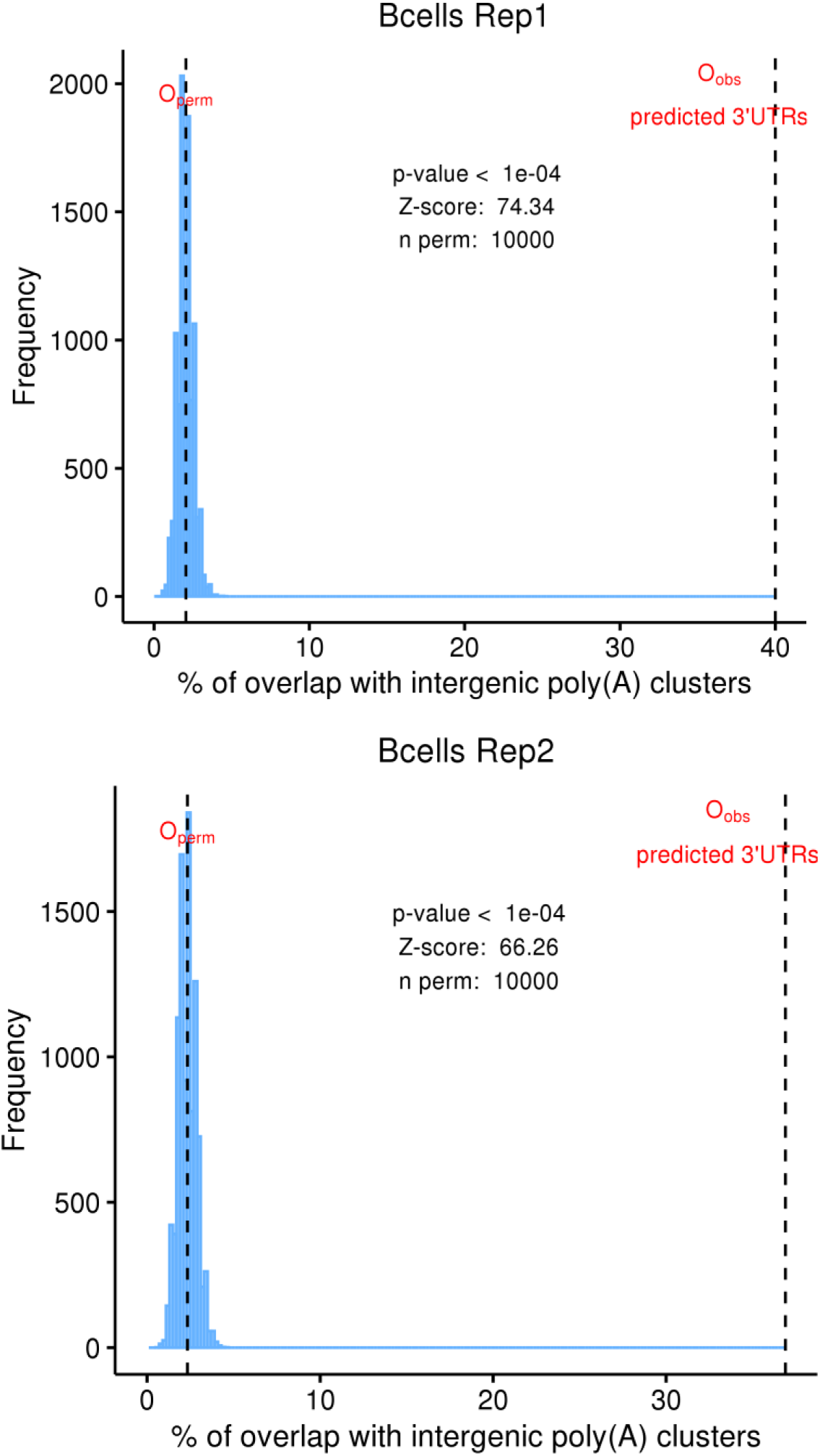
Overlap between randomly selected intergenic ERs and poly(A) sites. Distribution of overlap between randomly selected intergenic ERs and poly(A) sites from 10,000 permutations. O_perm_: mean overlap of the permuted distribution; O_obs_: observed overlap of 3’UTR predictions.

**Supplementary Figure S6.**
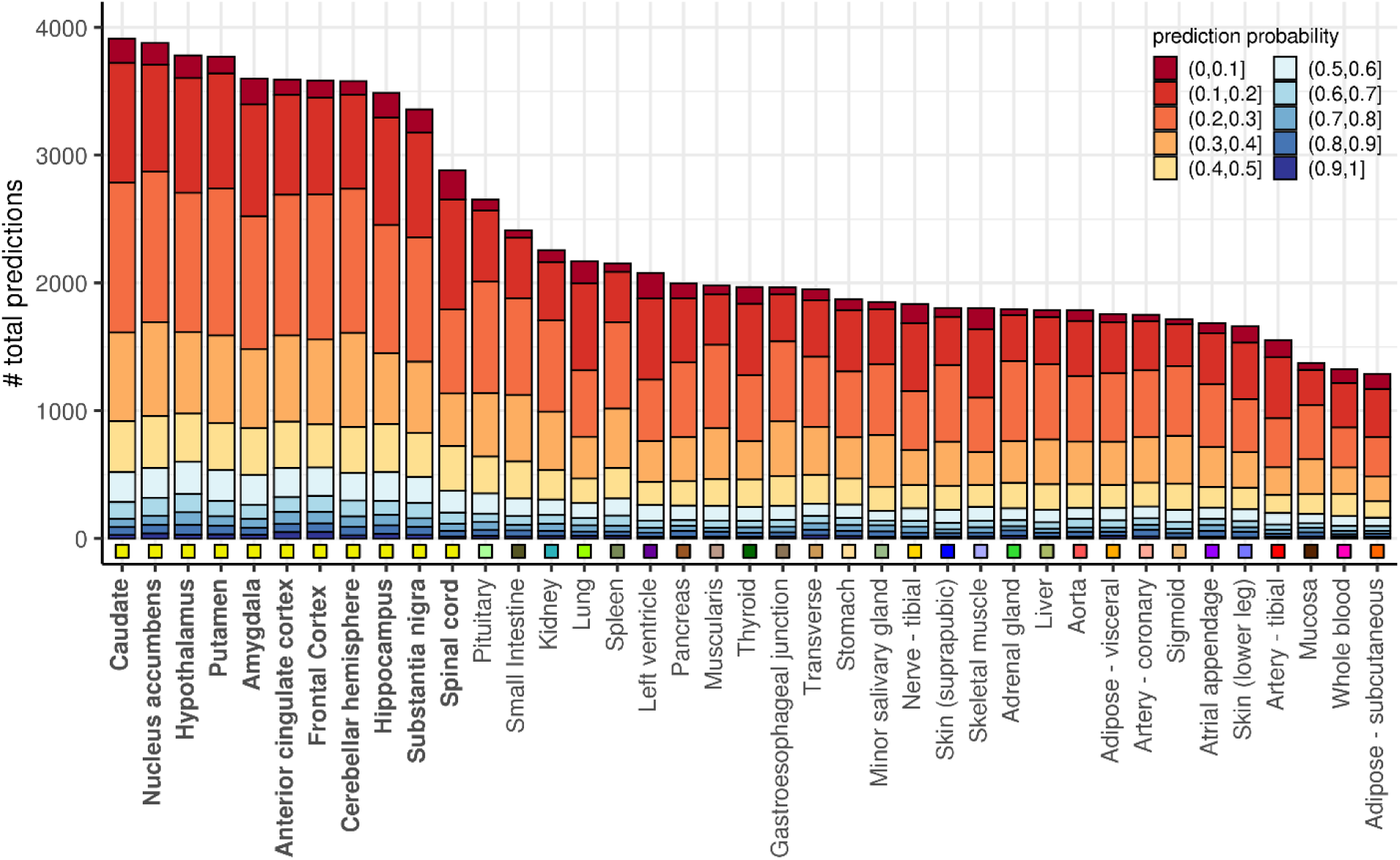
F3UTER predictions across 39 GTEx tissues. Bar plot showing the number of predictions in each tissue, grouped and color-coded according to their prediction probability scores. Tissues are sorted in descending order of the total number of predictions in each tissue. The square boxes below the bars are color-coded to group the tissues according to their physiology.

**Supplementary Figure S7.**
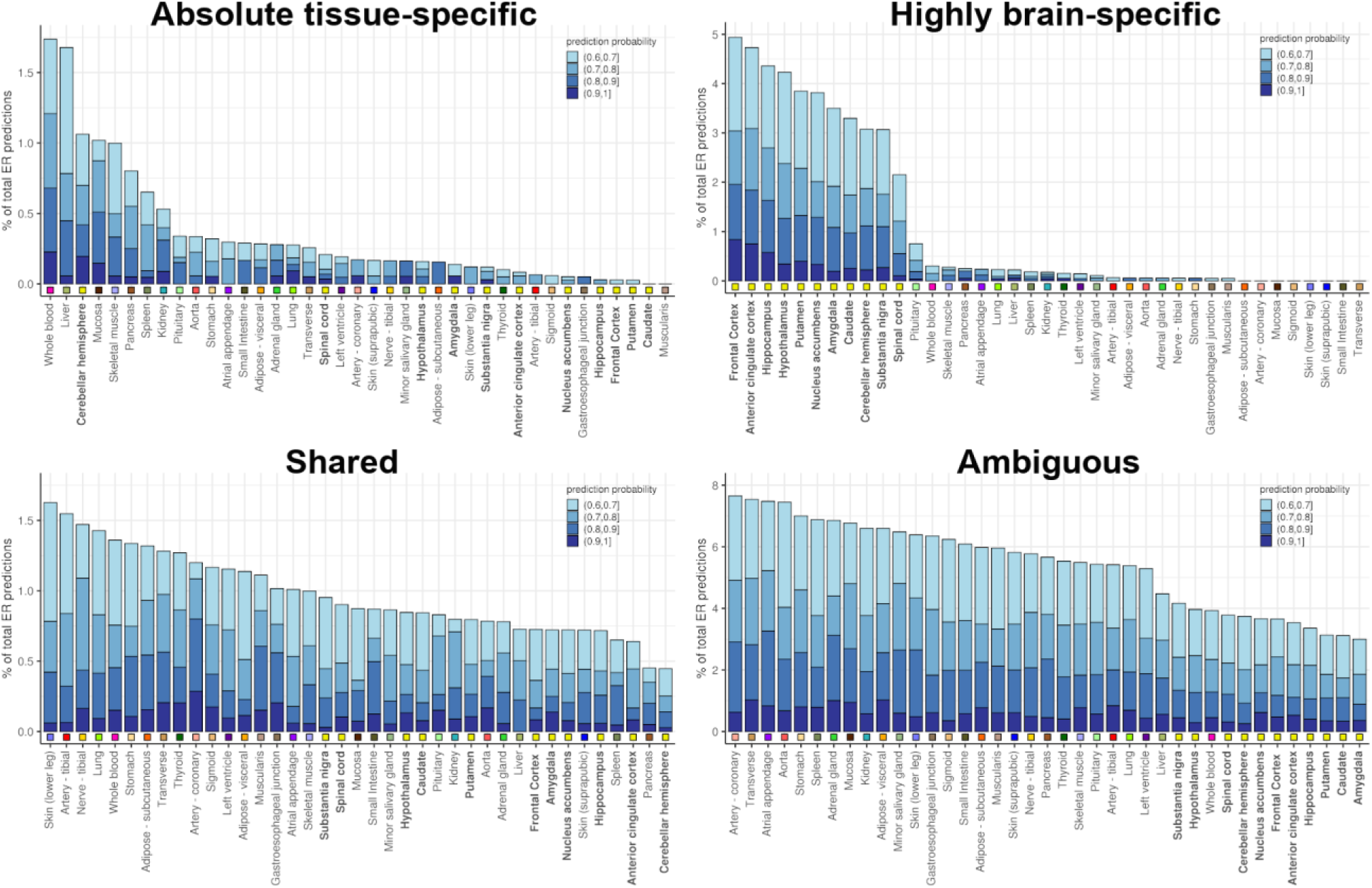
Categorisation of F3UTER predictions based on tissue-specificity. Bar plots showing the number of predictions grouped according to their tissue specificity across 39 tissues. Tissues are sorted in descending order of the number of predictions. The square boxes below the bars are color-coded to group the tissues according to their physiology.

**Supplementary Figure S8.**
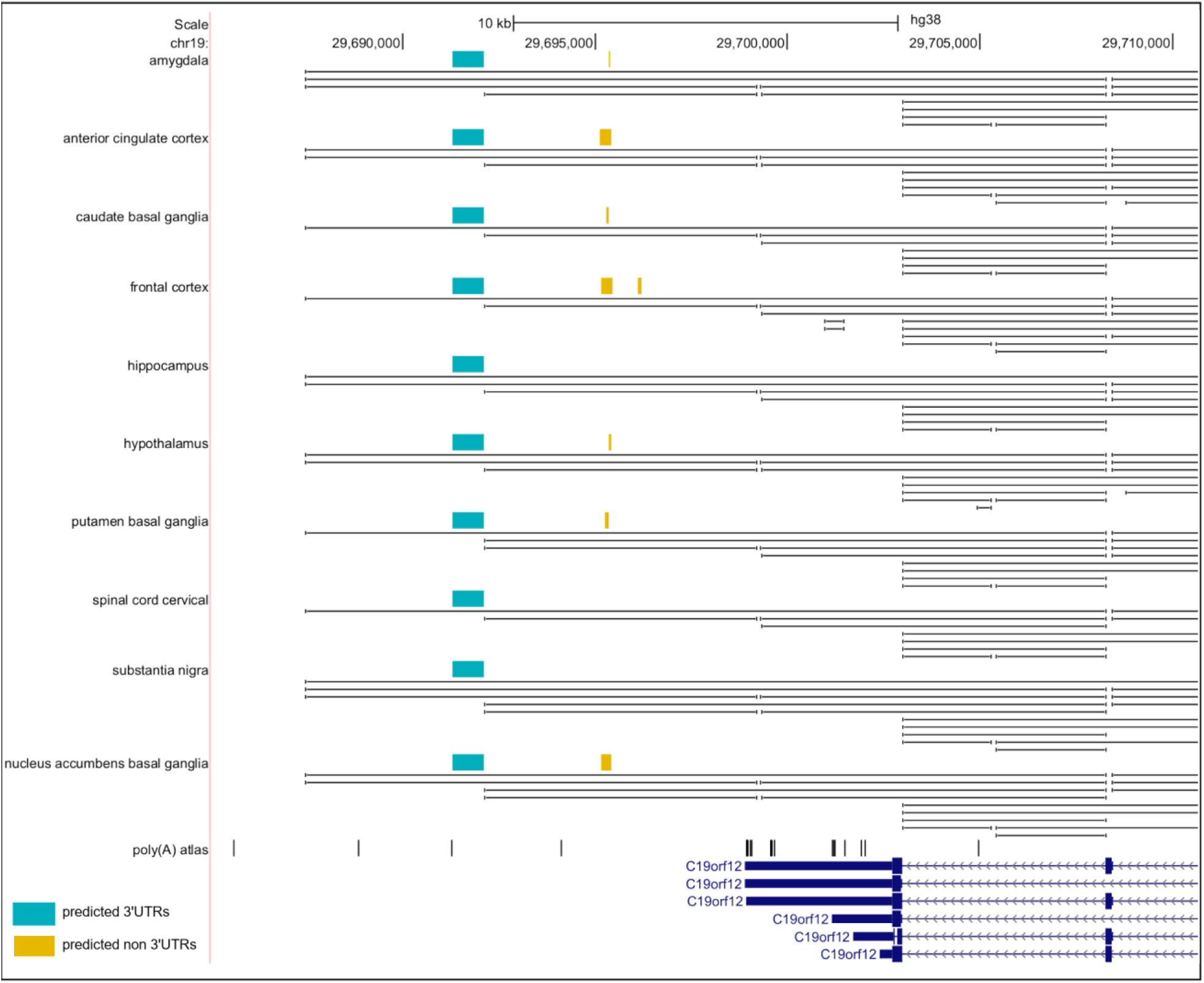
Unannotated 3’UTR associated with *C19orf12* in brain. Genomic view of the *C19orf12* locus displaying intergenic ERs and poly(A) sites from poly(A) atlas in the region. Two tracks are displayed for each tissue - the top track shows coloured boxes which represent the intergenic ERs, while the bottom track shows black lines which represent RNA-seq junction reads.

**Supplementary Figure S9.**
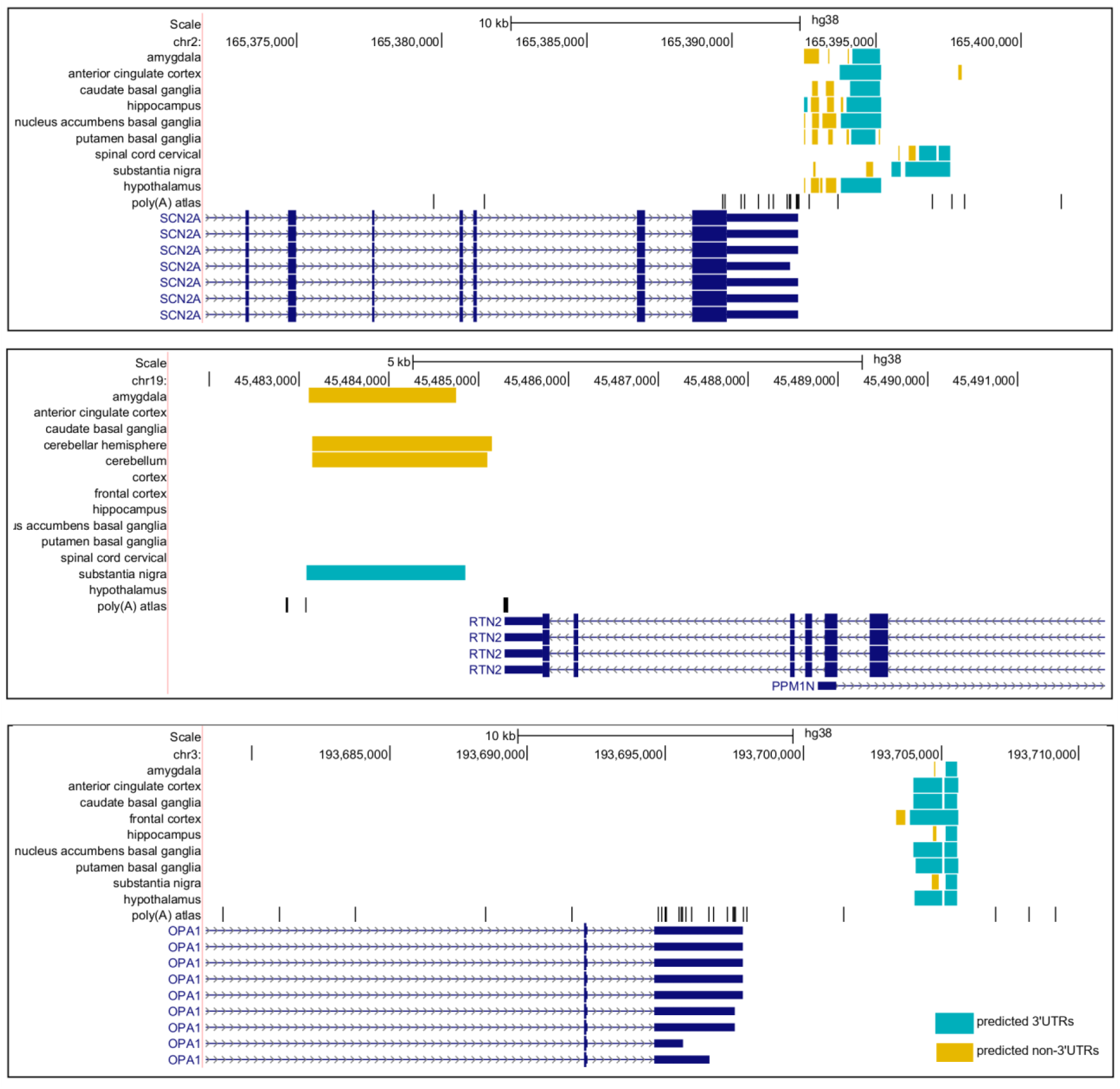
Examples of highly brain-specific unannotated 3’UTRs. Genomic view of genes (top: *SCN2A*; middle: *RTN2*; bottom: *OPA1*) associated with an unannotated 3’UTR in brain, displaying intergenic ERs and poly(A) sites from poly(A) atlas in the region.

**Supplementary Table 1.** List of RBPs with significantly enriched binding in the brain-specific unannotated 3’UTRs compared to shared unannotated 3’UTRs (*adjusted p* < 10^−5^). The enrichment p-value of the motifs were adjusted for multiple tests using a Bonferroni correction.

| Rank | RBP Name | RBP Ensembl id | p-value | Adjusted p-value |
| --- | --- | --- | --- | --- |
| 1 | *RBFOX1* | ENSG00000078328 | 5.83E-21 | 1.78E-18 |
| 2 | *KHSRP* | ENSG00000088247 | 4.12E-17 | 8.21E-15 |
| 3 | *ERI1* | ENSG00000104626 | 1.07E-15 | 3.85E-13 |
| 4 | *TIAL1* | ENSG00000151923 | 5.24E-15 | 2.41E-12 |
| 5 | *ELAVL3* | ENSG00000196361 | 6.95E-15 | 3.60E-12 |
| 6 | *CELF1* | ENSG00000149187 | 1.18E-12 | 1.16E-10 |
| 7 | *SSB* | ENSG00000138385 | 3.96E-13 | 1.77E-10 |
| 8 | *TARDBP* | ENSG00000120948 | 2.72E-12 | 5.09E-10 |
| 9 | *PUM2* | ENSG00000055917 | 1.55E-11 | 3.49E-09 |
| 10 | *ZFP36L2* | ENSG00000152518 | 7.52E-12 | 4.30E-09 |
| 11 | *ZFP36* | ENSG00000128016 | 7.52E-12 | 4.30E-09 |
| 12 | *HNRNPDL* | ENSG00000152795 | 7.37E-11 | 9.14E-09 |
| 13 | *AGO2* | ENSG00000123908 | 6.50E-10 | 1.11E-07 |
| 14 | *SRSF10* | ENSG00000188529 | 6.02E-10 | 1.58E-07 |
| 15 | *HNRNPAB* | ENSG00000197451 | 6.02E-10 | 1.58E-07 |
| 16 | *RBM5* | ENSG00000003756 | 3.52E-10 | 1.60E-07 |
| 17 | *HNRNPA2B1* | ENSG00000122566 | 4.09E-09 | 3.68E-07 |
| 18 | *ZRANB2* | ENSG00000132485 | 2.72E-09 | 5.80E-07 |
| 19 | *SRSF3* | ENSG00000112081 | 8.68E-09 | 1.49E-06 |
| 20 | *TRA2B* | ENSG00000136527 | 8.68E-09 | 1.75E-06 |
| 21 | *HNRNPD* | ENSG00000138668 | 8.12E-08 | 3.51E-05 |
| 22 | *AKAP1* | ENSG00000121057 | 5.94E-07 | 9.51E-05 |

